# Anti-HIV activity prediction and enrichment identify novel human targets for Anti-HIV therapy

**DOI:** 10.1101/159236

**Authors:** Shao-Xing Dai, Wen-Xing Li, Hui-Juan Li, Jia-Qian Liu, Jun-Juan Zheng, Qian Wang, Bi-Wen Chen, Yue-Dong Gao, Gong-Hua Li, Jing-Fei Huang

## Abstract

Human immunodeficiency virus (HIV) relies heavily on the host proteins to facilitate its entry and replication. Currently, more than 4000 human proteins are recorded to be involved in the HIV-1 life cycle. Identifying appropriate anti-HIV targets from so many host proteins is crucial to anti-HIV drug development, but a challenging work. Here we combined anti-HIV activity prediction and enrichment analysis to identify novel human targets for anti-HIV therapy. We firstly developed an accurate prediction tool named Anti-HIV-Predictor (AUC>0.96) to predict the anti-HIV activity of given compounds. Using this tool, we predicted 10488 anti-HIV compounds from ChEMBLdb. Then, based on this result and relationships of targets and compounds, we inferred 73 anti-HIV targets that enriched with anti-HIV compounds. The functional annotation and network analysis revealed that they directly or indirectly interact with 20 HIV proteins through neuropeptide signaling, GPCR signaling, cell surface signaling pathway, and so on. Nearly half of these targets overlap with the NCBI HIV dataset. However, the percentage of known therapeutic targets in these targets is significantly higher than that in the NCBI HIV dataset. After a series of feature analysis, we identified 13 novel human targets with high potential as anti-HIV targets, the inhibitors of which have experimentally confirmed anti-HIV activity. It is noteworthy that the inhibitors of REN and CALCA have better anti-HIV activity than CCR5 inhibitors. Taken together, our findings provide novel human targets for the host-oriented anti-HIV drug development and should significantly advance current anti-HIV research.

## Introduction

Acquired immune deficiency syndrome (AIDS) is a serious infectious disease caused by human immunodeficiency virus (HIV) infection^1^. As a chronic infectious disease that can directly impair human immune system, AIDS has become one of the world's most serious diseases ^2^. There are an estimated 39 million deaths caused by AIDS since its first recognition ^3^. According to the UNAIDS data, nearly 37 million people worldwide lived with HIV at the end of 2014. About 1.2 million people died from HIV-related causes globally in 2014. Therefore, prevention and treatment of HIV/AIDS are of great significance for human health and social stability.

Many scientists around the world are dedicated to finding scientifically proven strategies for HIV prevention and treatment. The discovery of new anti-HIV drugs is the main way to treat and control AIDS ^4^. Since the 1990s, with the continuous deepening of virus research and technological advances in drug discovery, a lot of anti-viral drugs were discovered, especially anti-HIV drugs ^2,4,5^ Many international pharmaceutical companies, such as Pfizer, Merck, GlaxoSmithKline and Roche, had invested heavily in anti-HIV drug development. Since the first anti-HIV drug, zidovudine, was approved by the US Food and Drug Administration (FDA) in 1987, there are more than thirty anti-HIV drugs approved by FDA ^6^. These drugs act mainly on reverse transcriptase, protease, integrase, Gp41 and so on ^7^. These anti-HIV drugs can effectively inhibit HIV replication and slow the progress of AIDS, thus greatly improve the life quality and survival time of AIDS patients. However, since these drugs mainly target a limited number of viral proteins, HIV can easily evade the selective pressure exerted by the drug through frequent mutations ^2^. With the widespread use of these drugs, it will result in cross-resistance and serious side effects. Therefore, the clinical application of these drugs is very limited ^5,8^. In addition, the discovery of anti-HIV target is greatly confined to a very small number of proteins encoded by the HIV genome.

HIV is an obligate intracellular pathogen and relies heavily on the host proteins for entry, replication and transmission. Therefore, we can achieve the goal of anti-HIV through targeting host proteins that are essential for HIV life cycle ^8, 9^. According to the Integrity database, multiple pharmaceutical companies have already launched the development of anti-HIV drug based on host protein. At present, a dozen of human proteins are used for anti-HIV drug development. These proteins are CCR5, CXCR4, CCR2B, HDAC, PKC, PXR, TLR3, ADH, CD4, TLR9, TSPO, and so on. Among them, as an antagonist of CCR5, maraviroc was approved for treatment of HIV infection in 2007. In view of the existing drug resistance, side effects, limited targets and many other issues, host-targeting anti-HIV has become an important strategy for the current anti-HIV drug development ^10-14^.

In recent years, many studies discovered a large number of HIV-associated host proteins and viral-host protein interactions by proteomics, RNA interference (RNAi), CRISPR screen, microarrays, RNA-Seq ^15-19^. These studies not only provide a solid foundation for revealing HIV infection mechanism at the level of viral-host molecular networks, they are also important for host-targeting anti-HIV drug development. However, there is little agreement among different studies and few human proteins have been validated ^17, 19^. Furthermore, the proteins identified by these high-throughput screening methods may not be targeted by drug because of lacking druggability ^20-22^. Currently, more than 4000 human proteins involved in the HIV-1 life cycle are documented in the HIV-1 Human Interaction Database ^23^. Identifying appropriate anti-HIV targets from so many host proteins is crucial to anti-HIV drug development, but a challenging work.

Therefore, we aimed to identify novel human targets for anti-HIV therapy by anti-HIV activity prediction and enrichment (APE) method. In our previous study, we first used the APE method to infer anti-cancer targets and identified HKDC1 as a novel potential therapeutic target for cancer ^24^. In this study, the APE method mainly contains two steps as shown in **Figure 1**: anti-HIV activity prediction and enrichment analysis of anti-HIV activity. We firstly developed Anti-HIV-Predictor to predict the anti-HIV activity of a given compound by integrating three rapid and accurate computational methods. Then the anti-HIV activities of all compounds with human targets in ChEMBLdb were predicted using Anti-HIV-Predictor. In the second step, we performed a hypergeometric test to determine the targets that significantly enriched with anti-HIV compounds. Using this strategy, we identified 73 potential human targets for anti-HIV therapy. These human targets directly or indirectly interact with HIV proteins through neuropeptide signaling, GPCR signaling, cell surface signaling pathway, and so on. Nearly half of these targets overlap with the NCBI HIV dataset. However, the percentage of known therapeutic targets in these targets is significantly higher than that in the NCBI HIV dataset. In addition, 37 of these targets belong to G-protein coupled receptor family, which indicates that the GPCRs may be more important for HIV life cycle than ever known. After a series of feature analysis, we identified 13 novel human targets with high potential as anti-HIV targets. The inhibitors of the 13 novel targets have experimentally confirmed anti-HIV activity. The results of this study provide important ideas and guidance for the host-oriented anti-HIV drug development.

**Figure 1.**
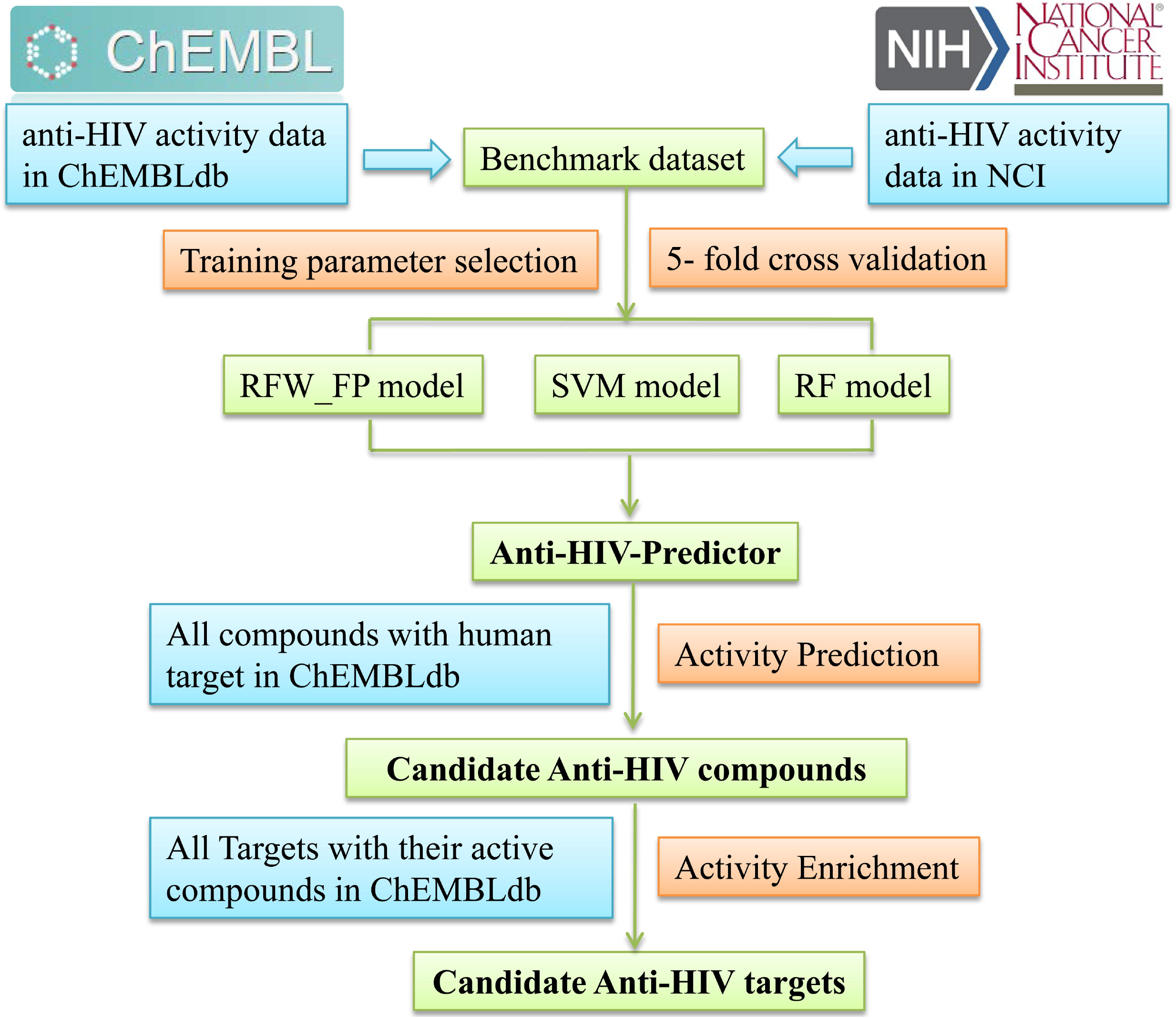
The flowchart of identifying novel human targets for anti-HIV therapy. After construction of benchmark dataset, three models (RFW_FP model, SVM model and RF model) were generated to predict anti-HIV activity by training, parameter selection and 5-fold cross validation. The three models were incorporated to establish Anti-HIV-Predictor. Then Anti-HIV-Predictor was used to predict the anti-HIV activities of all compounds with human targets in ChEMBLdb. Finally, the targets enriched with anti-HIV compounds were inferred as candidate anti-HIV targets.

## Results

### Development and performance of Anti-HIV-Predictor

Anti-HIV-Predictor was developed to predict the anti-HIV activity of given compounds (**Figure 1**). Anti-HIV-Predictor firstly integrated all the data of anti-HIV activity from ChEMBL and NCI database to construct a benchmark dataset. Then, using the benchmark dataset, three prediction models were generated by training, parameter selection and validation. The three models are relative frequency weighted fingerprint (RFW_FP) based model, support vector machine (SVM) model and random forest (RF) models, respectively. Last, three models (RFW_FP model, SVM model and RF model) were incorporated to predict the anti-HIV activity of chemical compounds. The details for development of Anti-HIV-Predictor are given in the **Materials and methods** section.

The classification performance of the models was assessed in terms of accuracy, precision, recall and F1 score (**Figure 2**). As 10 runs of 5-fold cross-validation method were used, these scores were averaged. Over the 10 runs, their standard deviations were also reported. As shown in **Figure 2**, the RFW_FP model obtains the statistical average of 93%, 87%, 90%, and 88% for accuracy, precision, recall, and F1 score, respectively. The accuracy, precision, recall, and F1 score of SVM model are 96%, 95%, 91%, and 93%, respectively. RF model performs best with accuracy of 96% and precision of 99%. The overall performance of the RFW_FP, SVM and RF models was also quantified by receiver operating characteristic curve (ROC). For each model, the area under the curve (AUC) was calculated (**Figure 2**). The AUC values of the RFW_FP, SVM and RF models are 0.96, 0.97 and 0.98, respectively. All three models achieve AUC value greater than 0.96, which reveals the excellent effectiveness of the models.

**Figure 2.**
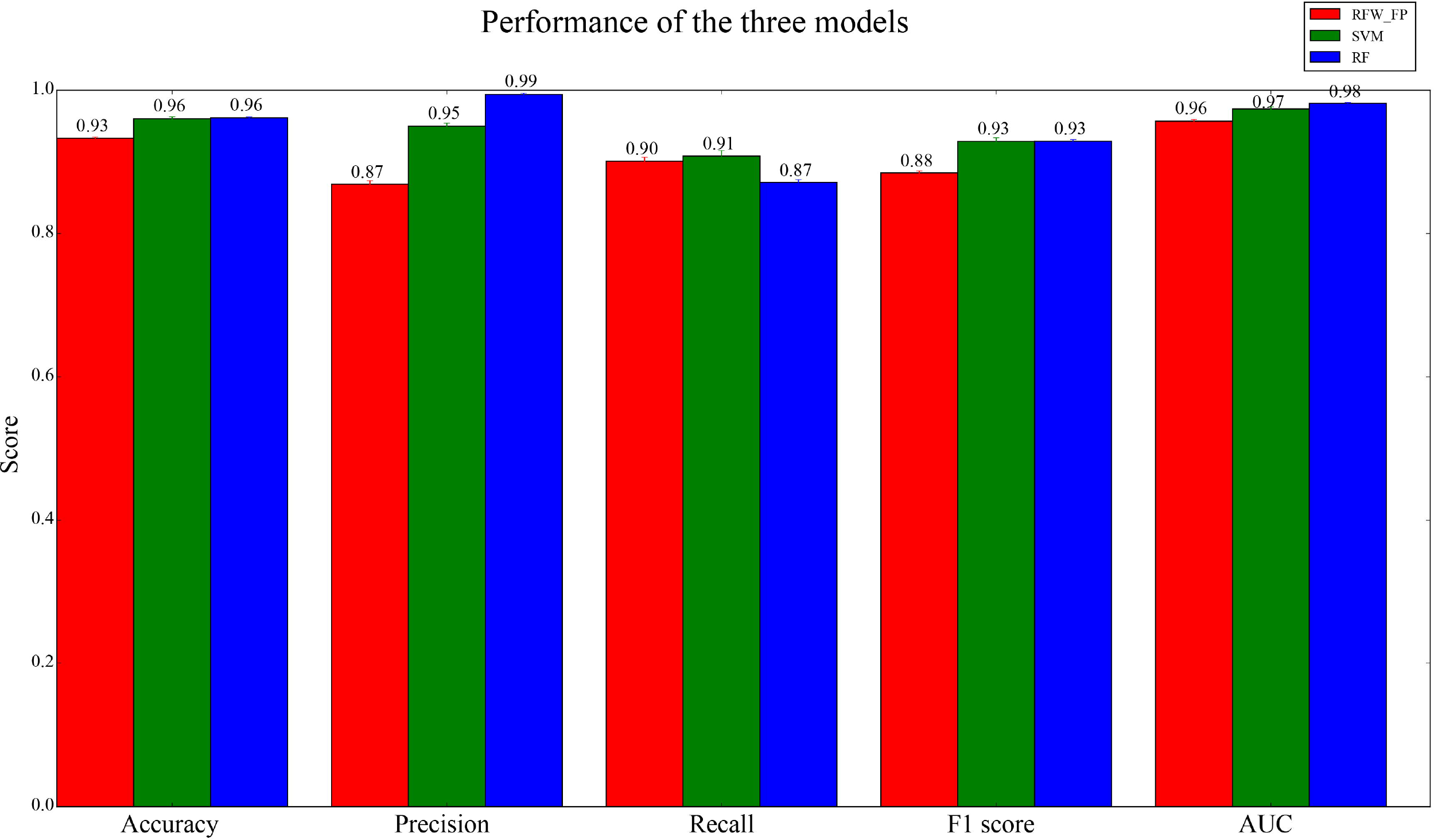
The performance of Anti-HIV-Predictor. The bar chart illustrates the statistical average results for 10 runs of 5-fold cross validation derived from the RFW_FP model (red), SVM model (green) and RF model (blue), respectively. The mean and standard deviation values of accuracy, precision, recall, F1 score and AUC were shown. Vertical lines indicate the standard deviations.

### The characteristic of relationship between human targets and their ligands in ChEMBLdb

In order to infer potential anti-HIV human target, all human targets were downloaded from ChEMBLdb. Their ligands whose activity is better than 10uM were also downloaded. Totally, we obtained 1924 targets and 251825 their active compounds. We characterized the relationship between human targets and their ligands (**Figure 3**). There are 384768 target-compound pairs for the 1924 targets and their active compounds. 72% (1377) of these targets have no more than 100 ligands. With the number of their ligands increases, the number of targets decreases drastically (**Figure 3A**). There are only 11% (209) targets having more than 500 ligands. All the compounds were grouped according to their target types (**Figure 3B**). 52.2% (200194) compounds are enzyme inhibitors. 30.6% (117509) compounds are membrane receptor modulator. In addition, these compounds can also act on ion channels, transcription factors, transporters, epigenetic regulators and other proteins.

**Figure 3.**
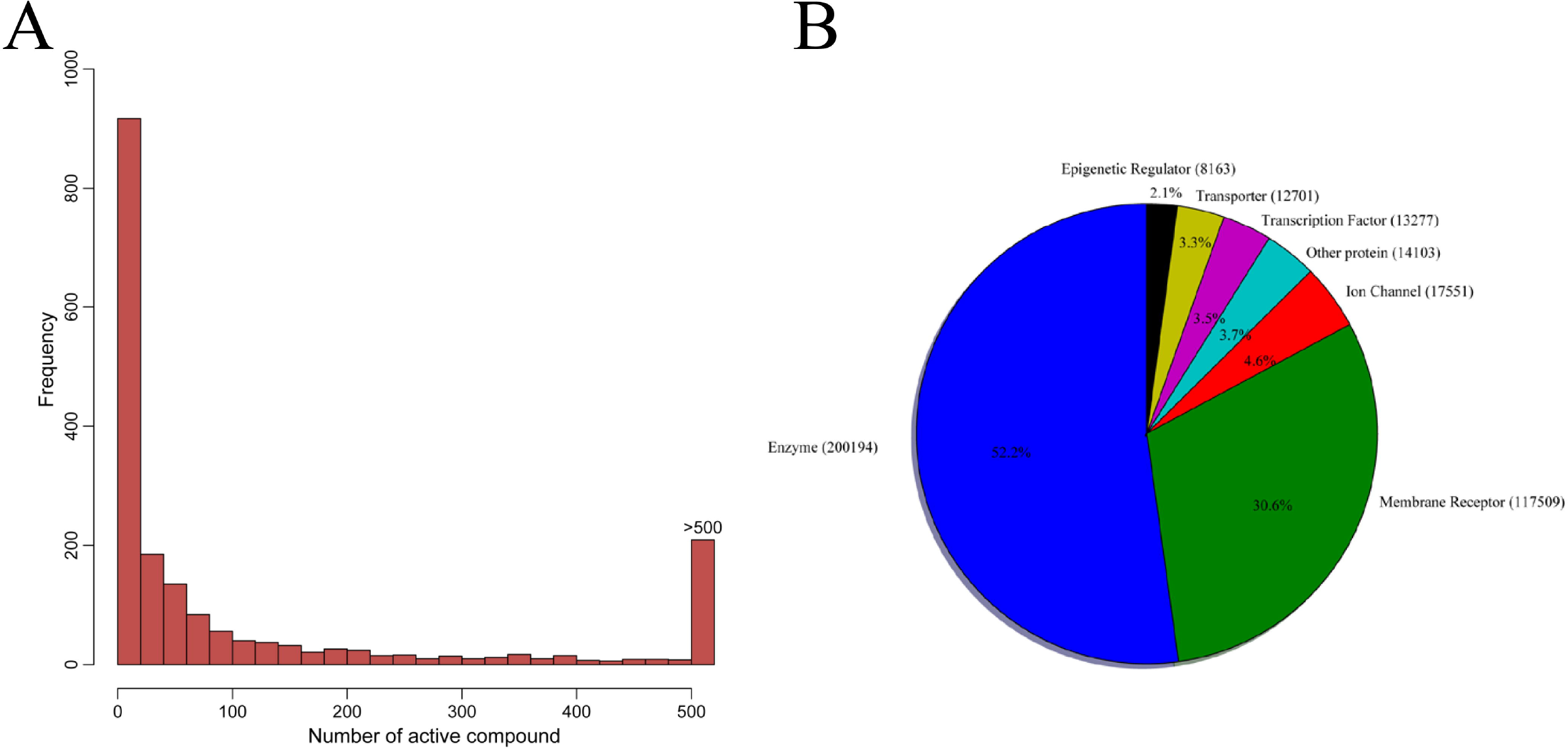
The statistics of human targets and their ligands in ChEMBLdb. **A)**. Histogram plot of the frequency of the targets with different numbers of compounds. With the number of their ligands increases, the number of targets decreases drastically. **B)** Pie chart displaying numerical proportion of different classes of all the compounds. All the compounds were classified by their target types.

### Prediction of anti-HIV activity for all compounds with human target in ChEMBLdb

To infer anti-HIV human targets by anti-HIV activity enrichment, we predicted the anti-HIV activity of the 251825 compounds that act on human proteins using Anti-HIV-Predictor. The results of the computational screen are shown in **Figure 4**. The inactive compounds were shown as blue dots (RFW_TC P-value≥0.05, SVM probability and RF probability≤0.5). The green dots represent the compounds with anti-HIV activity supported by one or two models (RFW_TC P-value < 0.05 or SVM probability > 0.5 or RF probability > 0.5). The red dots represent the compounds with anti-HIV activity supported by all three models (RFW_TC P-value < 0.05 and SVM probability > 0.5 and RF probability > 0.5). A total of 10488 compounds were predicted as anti-HIV compounds by all three models, which are accounting for 4% (10488/251825) of all compounds in the ChEMBLdb.

**Figure 4.**
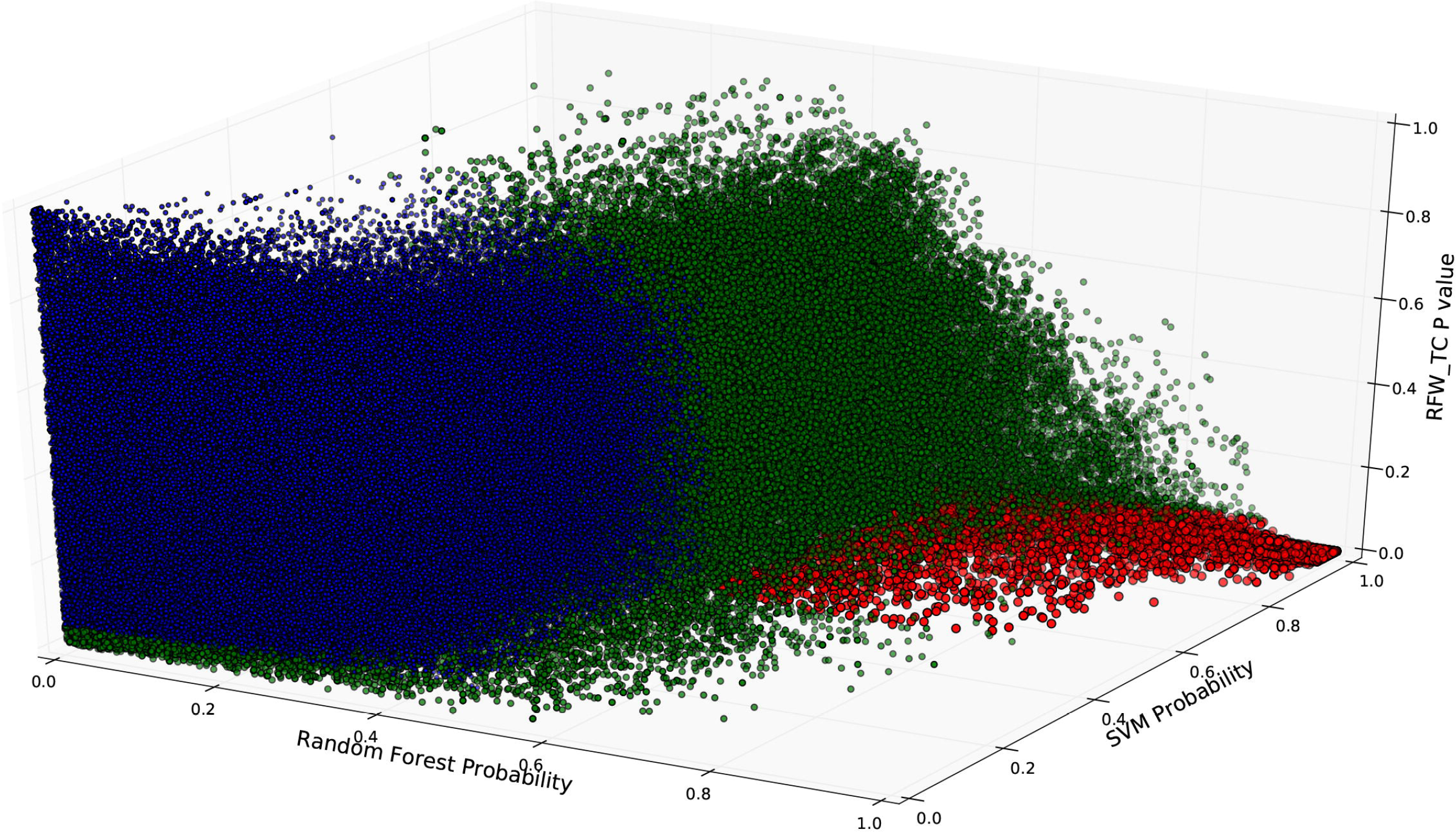
The predicted results for all compounds in ChEMBLdb using Anti-HIV-Predictor. The predicted scores for all compounds in ChEMBLdb were shown in the three-axis plot based on the three models (RFW_FP model, SVM model and RF model). Each dot represents a compound. The blue dot means the compound with no anti-HIV activity. The green dot means the compound with anti-HIV activity supported by one or two models. The red dot indicates the compound with anti-HIV activity predicted by all three models.

### Inferring 73 potential anti-HIV human targets

The identification of anti-HIV human targets is of great value for the development of anti-HIV drugs. We have predicted tens of thousands of compounds with anti-HIV activity above. Based on the result, we can infer potential anti-HIV targets that enriched with anti-HIV compounds. A total of 73 human targets (P_adj < 0.05) were inferred using the APE method (**Supplemental Table S1**). These targets play their functions in multiple biological processes such as neuropeptide signaling pathway, positive regulation of cytosolic calcium ion concentration, G-protein coupled receptor signaling pathway and cell surface receptor signaling pathway (**Figure 5A**). They perform a variety of functions, mainly including neuropeptide binding, peptide hormone binding, somatostatin receptor activity, peptide binding, melanocortin receptor activity and so on. These targets are in contact with each other closely through genetic interaction, pathway and physical interaction (**Figure 5B**). As shown in the network, these targets connect with 20 HIV proteins closely and extensively. It suggests these targets are very important and truly involved in the HIV life cycle.

**Figure 5.**
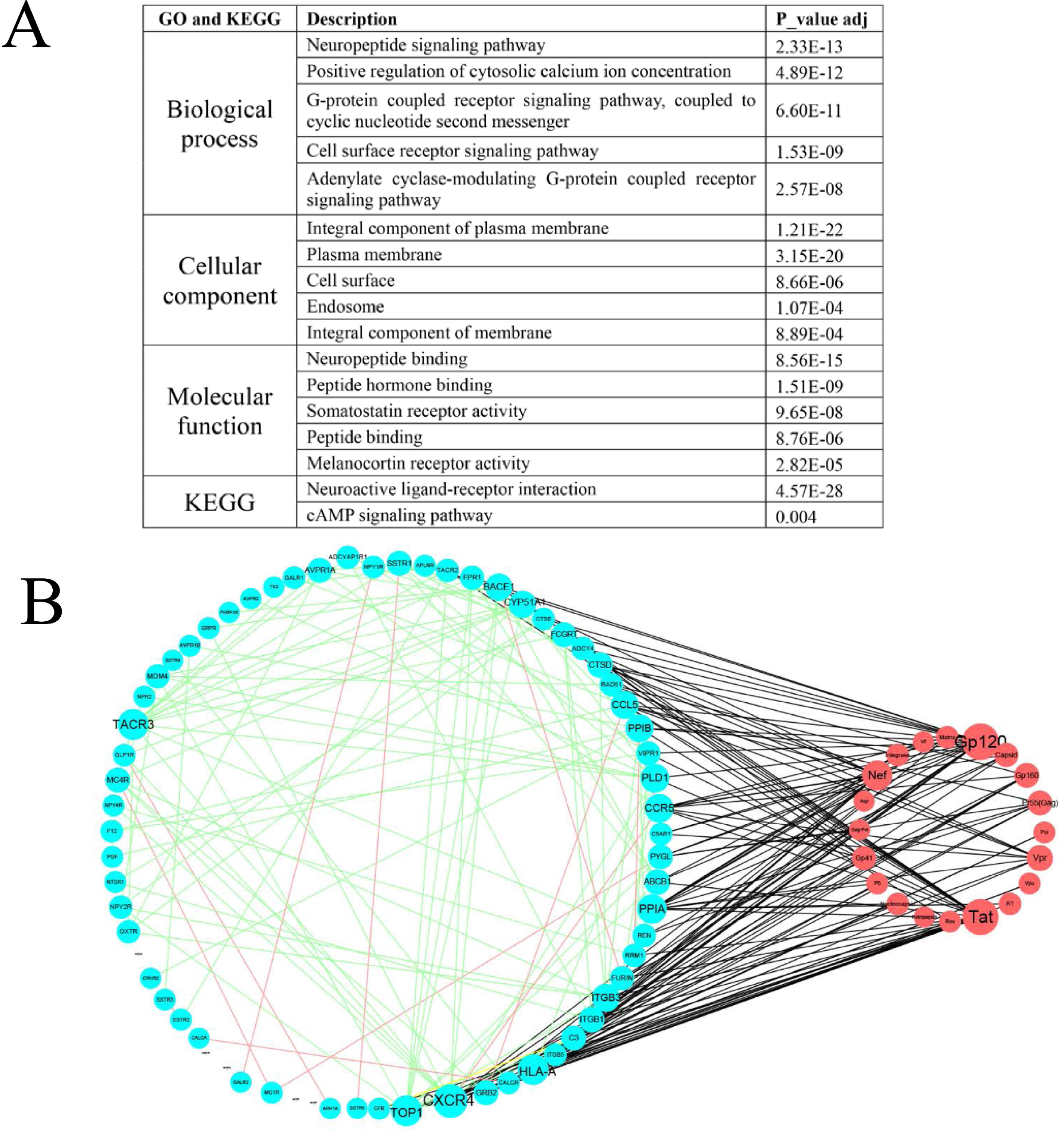
The functional annotation and network of the candidate anti-HIV targets. **A)** Geneontology and KEGG enrichment analysis of the candidate anti-HIV targets. Only the top five significantly enriched Go terms and pathways were shown. P_value_adj means Benjamini-adjusted P-value. **B)** The network of the candidate anti-HIV human targets and HIV proteins. The aqua and red and circles represent human targets and HIV proteins, respectively. The interactions between human target and HIV protein are shown as black lines. These targets are in contact with each other closely through genetic interaction, pathway and physical interaction, which are shown as green, yellow and red lines, respectively. The node size is proportional to the number of connections.

### Comparison of the predicted targets in this study with NCBI HIV dataset

In order to compare our study with other studies, we downloaded all HIV-related host factors from HIV-1 interactions database in NCBI (**Supplemental Table S3**) ^23^. These host factors and HIV proteins affect each other through protein interaction and replication interaction. Protein interaction and replication interaction involved in 3649 and 1325 human proteins, respectively. Close half (32/73) of targets identified in this study overlap with the NCBI HIV dataset (**Figure 6A**). In particular, most of the overlapped proteins participate in protein interaction with HIV. Next, we compared the percentages of known therapeutic targets between our study and the NCBI HIV dataset (**Figure 6B**). We found 32.9% of these targets (24 of 73) are approved targets. 75.3% of these targets (55 of 73) are used in clinical trials. The percentage of known targets (approved or clinical targets) within these 73 targets is four times than that within the NCBI HIV dataset. These results indicated that the anti-HIV compounds-enriched targets have high likelihoods of being developed into clinical targets (>75%). Thus, these 73 targets represent potential anti-HIV human targets that should be validated further using other methods.

**Figure 6.**
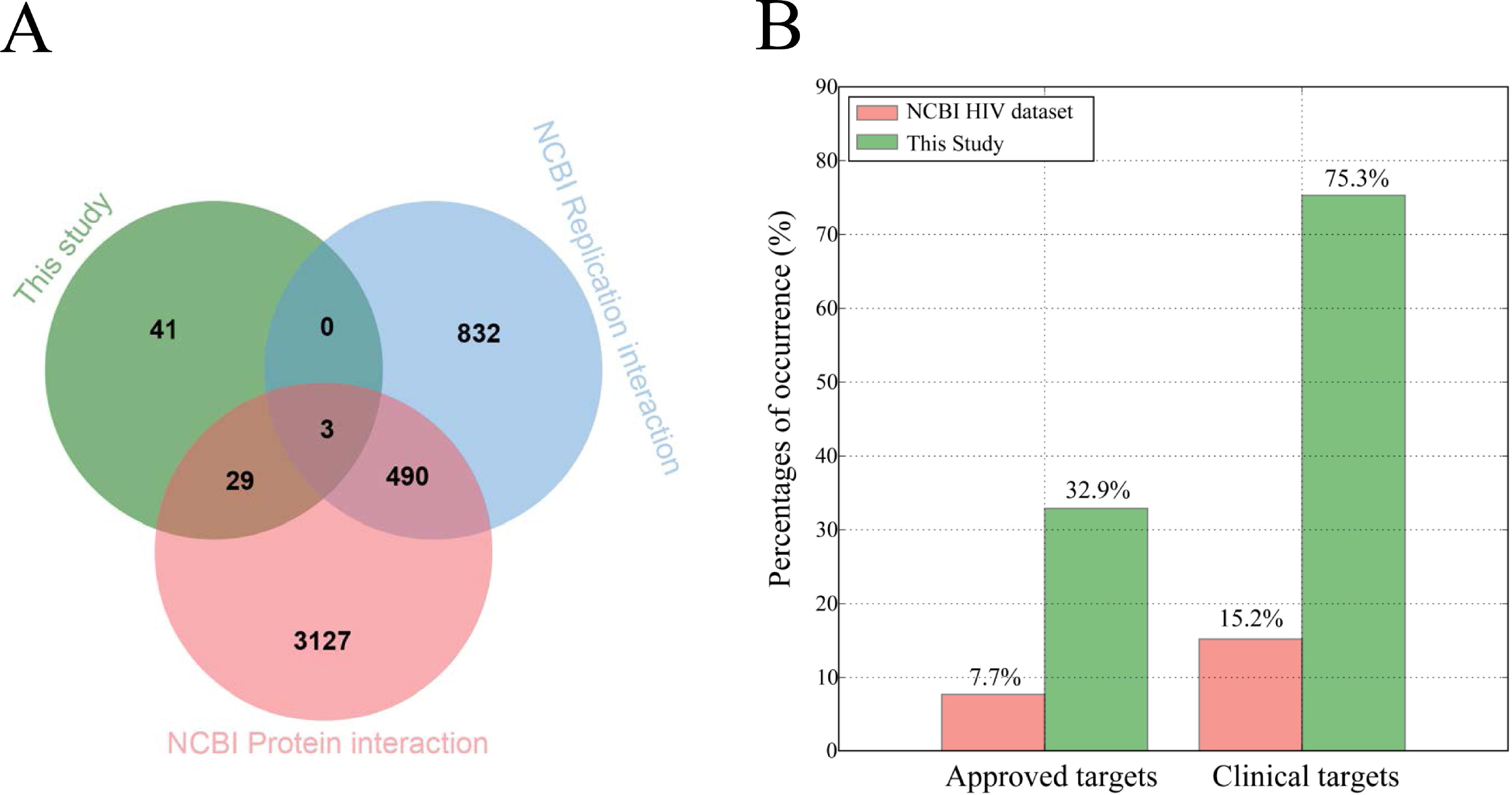
Comparison of our study with NCBI HIV dataset. **A)** Venn diagram showing the overlap between this study and the NCBI HIV dataset. The NCBI HIV dataset, obtained from HIV-1 Human Interaction Database, contains HIV-1 protein-human protein interactions (NCBI Protein interaction) and interaction effects upon HIV-1 replication (NCBI Replication interaction). **B)** Comparison of the percentages of known therapeutic targets between this study and the NCBI HIV dataset. The percentage of known targets (approved or clinical trial targets) within these targets of this study is four times than that within the NCBI HIV dataset.

### Identifying 13 novel human targets whose inhibitors have potent anti-HIV activity

To further evaluate the anti-HIV potential of the 73 targets, we checked whether the inhibitors of these targets have experimentally confirmed anti-HIV activity. We found 16 human targets whose inhibitors have anti-HIV activity through mapped the inhibitors into anti-HIV activity data from ChEMBLdb and NCI (**Figure 7** **and** **Table 1**). Among them, CCR5, CXCR4 and ABCB1 are currently already used for anti-HIV therapy. The other 13 targets, highlighted with underline in **Table 1**, represent novel human targets identified in this study. Of the 13 novel targets, 4 targets have approved drugs for other diseases, 11 targets were used in clinical trials for other diseases (**Table 1**). Furthermore, 10 of the 13 novel targets are closely related to the HIV life cycle. These features indicate the good druggability of these novel targets. For most of these novel targets, the anti-HIV activities of their inhibitors are about 1 uM, which is at the same level of activity compared with known anti-HIV targets (**Figure 7**). It is noteworthy that the inhibitors of REN and CALCA have better anti-HIV activity (0.136 uM, 0.102 uM, respectively) than CCR5 inhibitors (0.227 uM). Furthermore, the compound CHEMBL470508, the inhibitor of CTSD, show potent anti-HIV activity (0.008uM) than the approved anti-HIV drug maraviroc (0.016uM). Therefore, the 13 novel human targets identified in this study have high potential as the targets for HIV therapy.

**Figure 7.**
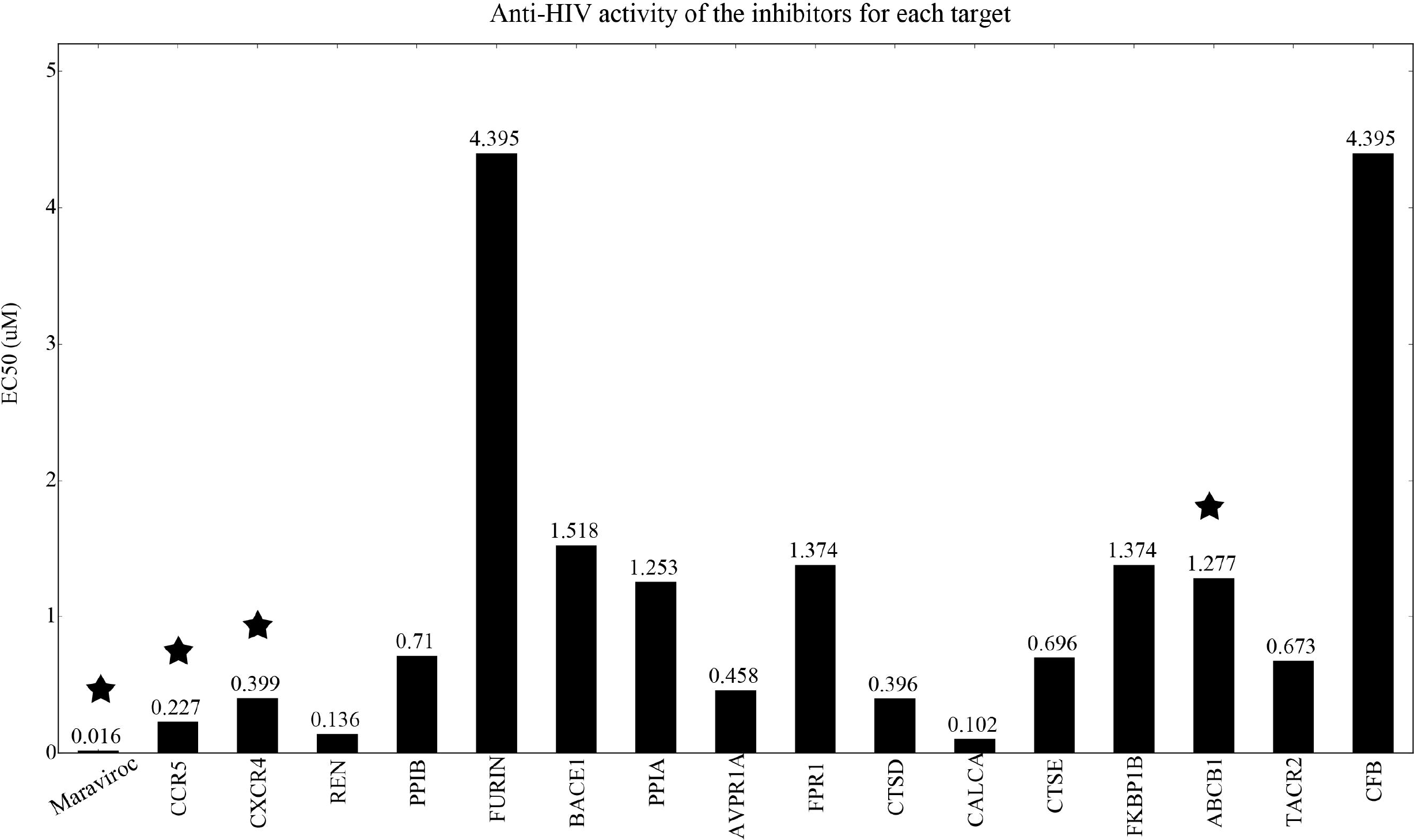
The bar chart displaying anti-HIV activity of the inhibitors for 13 novel human targets and known anti-HIV targets. The known anti-HIV targets and drug maraviroc are marked with asterisks. The other targets are identified as novel human targets in this study. All the activity values (uM) were labeled on the top of bar.

**Table 1.** The 13 novel human targets identified in this study compared with currently used anti-HIV targets.

| Gene | Approved Target? | Clinical Target? | <sup>c</sup> Related to HIV? | Anti-HIV Target? | <sup>d</sup> Anti-HIV activity of inhibitors (EC50, uM) | Adjusted P-value |
| --- | --- | --- | --- | --- | --- | --- |
| <sup>a</sup> **CCR5** | √ | √ | √ | √ | 0.227 | 0 |
| **Protease** | √ | √ | √ | √ | 0.557 | 0 |
| **RT** | √ | √ | √ | √ | 0.047 | 0 |
| **Integrase** | √ | √ | √ | √ | 0.364 | 1.91E-281 |
| **CXCR4** | √ | √ | √ | √ | 0.399 | 1.19E-167 |
| <u><sup>b</sup> REN</u> | √ | √ | √ | × | 0.136 | 2.16E-155 |
| <u>PPIB</u> | √ | √ | √ | × | 0.71 | 9.39E-24 |
| <u>FURIN</u> | × | √ | √ | × | 4.395 | 2.42E-21 |
| <u>BACE1</u> | × | √ | √ | × | 1.518 | 5.24E-15 |
| <u>PPIA</u> | √ | √ | √ | × | 1.253 | 7.75E-14 |
| <u>AVPR1A</u> | √ | √ | Unknown | × | 0.458 | 1.42E-10 |
| <u>FPR1</u> | × | √ | √ | × | 1.374 | 1.79E-08 |
| <u>CTSD</u> | × | √ | √ | × | 0.396 | 1.99E-08 |
| <u>CALCA</u> | × | √ | Unknown | × | 0.102 | 4.16E-05 |
| <u>CTSE</u> | × | × | √ | × | 0.696 | 6.03E-05 |
| <u>FKBP1B</u> | × | × | Unknown | × | 1.374 | 2.40E-04 |
| **ABCB1** | × | √ | √ | √ | 1.277 | 4.33E-04 |
| <u>TACR2</u> | × | √ | √ | × | 0.673 | 5.19E-04 |
| **GP41** | √ | √ | √ | √ | 1.128 | 1.21E-02 |
| <u>CFB</u> | × | √ | √ | × | 4.395 | 2.63E-02 |

**Note:** **a.** The seven proteins CCR5, CXCR4, ABCB1, Protease, RT, Integrase and GP41, shown with bold fonts, are currently used as the targets for anti-HIV therapy. Of them, protease, RT, integrase, and GP41 are HIV proteins. **b.** These proteins identified as novel human targets for anti-HIV therapy were highlighted with underline. **c.** This column shows whether the proteins are related to HIV life cycle according to HIV-1 Interactions database. **d.** The column shows the average anti-HIV activity of the inhibitors for each target. For the HIV proteins protease, RT, integrase and GP41, the anti-HIV activity of their representative inhibitors nelfinavir, nevirapine, raltegravir, and enfuvirtide were shown, respectively.

## Discussion

The failure of 30 years of HIV vaccine development ^25, 26^, as well as the prevalence of drug-resistant HIV ^27-29^, emphasizes the need for new, effective and affordable anti-HIV drugs. New target discovery will promote discovery of new anti-HIV drugs. Therefore, identifying host proteins as novel anti-HIV targets is crucial to the development of anti-HIV drugs, especially host-targeting drugs. RNAi and CRISPR can be used to identify HIV-related host factors. However, there are some shortcomings for the two methods. RNAi-based screens have identified thousand candidate host factors, but there is little agreement among studies ^19^. In contrast, CRISPR screen only discovered five host factors, two of which are known factors CCR5 and CD4 ^19^. Therefore, new methods for identification of human target are needed to improve this situation. In this study, benefited from massive activity data between target and compound in ChEMBLdb and NCI, we can identify potential human targets for anti-HIV therapy using the APE method. Firstly, we predicted 10488 compounds with anti-HIV activity from 251825 compounds in ChEMBLdb using Anti-HIV-Predictor. Then we inferred 73 human targets significantly enriched with predicted anti-HIV compounds by enrichment analysis. Finally, we identified 13 novel human proteins with high potential as anti-HIV targets through checking confirmed anti-HIV activity. They are REN, PPIB, FURIN, BACE1, PPIA, AVPR1A, FPR1, CTSD, CALCA, CTSE, FKBP1B, TACR2 and CFB. It is worthy to study these targets further in detail. And strict experimental validation of these proteins as anti-HIV targets is needed.

The CCR5 antagonist, maraviroc, is approved for treatment of HIV infection by the FDA in 2007. The success of maraviroc indicates host-targeting anti-HIV is an important strategy for the deployment of anti-HIV drugs. CCR5, as a protein on the surface of white blood cells, is involved in the immune system as it acts as G protein–coupled receptor for chemokines ^30^. HIV initially uses CCR5 as co-receptor to enter and infect host cells ^31^. G protein-coupled receptors signaling pathway can be hijacked by HIV revealed by previous studies ^32^. The 73 human targets inferred by enrichment analysis are involved in G-protein coupled receptor signaling pathway (P-value=6.60E-11) and cell surface receptor signaling pathway (P-value=1.53E-9). Among them, 37 human targets belong to G-protein coupled receptor family (**Supplemental Table S1**). This result suggests the G protein-coupled receptor signaling pathway may be more important for HIV life cycle than ever known. Furthermore, these targets are significantly enriched in neuropeptide signaling pathway (P-value=2.3E-13). HIV infection will cause HIV-associated distal symmetric polyneuropathy and other HIV-related peripheral neuropathies. Our study indicates these targets may be involved in the neuropathy process induced by HIV infection.

We identified 13 novel human targets whose inhibitors have anti-HIV activity through mapped the inhibitors into anti-HIV activity data from ChEMBLdb and NCI (**Figure 7** **and** **Table 1**). The anti-HIV activities of the inhibitors for the 13 novel human targets were compared with CCR5. The average anti-HIV activity of the antagonists of CCR5 is 0.227 uM. The average anti-HIV activities of the inhibitors for the 13 novel human targets are ranging from 0.102 to 4.395 uM. For most of these novel targets, the anti-HIV activities of their inhibitors are about 1 uM, which is at the same level of activity compared with known anti-HIV targets (**Figure 7**). Especially, compound CHEMBL470508, the inhibitor of CTSD, show potent anti-HIV activity (0.008uM) than the approved anti-HIV drug maraviroc (0.016uM). These results suggest that some of the 13 novel human targets identified in this study may be served as important targets for HIV therapy like CCR5.

In summary, identifying appropriate anti-HIV targets from thousands of host proteins is a challenging work. Benefited from massive data of the target activities and anti-HIV activities, we developed the APE method to identify potential human targets for anti-HIV therapy. We identified 13 novel human targets after a series of steps, including development of Anti-HIV-Predictor, prediction of anti-HIV compounds, enrichment analysis of anti-HIV activity, and target features filtering. The method APE and the 13 novel anti-HIV human targets provided here will have an immediate effect on the development of new anti-HIV drugs, and should significantly advance current anti-HIV research.

## Materials and methods

### Construction of anti-HIV benchmark dataset and collection of information of targets and compounds

Anti-HIV activity data were downloaded from ChEMBLdb ^33,34^ and NCI ^35^. In ChEMBLdb, the compound whose target is "human immunodeficiency virus type 1" and with the activity better than 10µmol/L was considered as active compounds. In NCI, the compound with more than 2 replication experiments and with EC_50_ less than 10μmol/L was considered as active compounds. And the other compounds with EC_50_ more than 100μmol/L were considered as inactive compounds. Finally, all compounds in the two databases were integrated by removed the conflict and replicated compounds. This procedure yielded 9584 active and 23998 inactive compounds, respectively. The active and inactive datasets were used as benchmark datasets to generate models to predict the anti-HIV activity of chemical compounds. The detailed method of constructing benchmark dataset can be found in **Part 1 of Supplemental material**. To infer anti-HIV targets, we collected the information concerning the targets and their compounds from ChEMBLdb. The structures of all related compounds and information of human targets were downloaded.

### Development of Anti-HIV-Predictor

We incorporated three models (RFW_FP model, SVM model and RF model) as Anti-HIV-Predictor to predict the anti-HIV activity of chemical compounds. The RFW_FP model was generated as follows. Relative Frequency-Weighted Fingerprint (RFW_FP) was used in our previous study and powerful to distinguish the active and inactive compounds for anti-cancer ^24,36,37^. Firstly, RFW_FP was used to calculate the compound fingerprints as follows:

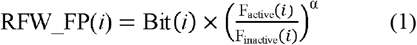

where *i* represents *i*th Daylight fingerprint. In Daylight theory, each compound contains more than one and less than 1024 fingerprints. RFW_FP(*i*) is *i*th relative frequency-weighted fingerprint. Bit(*i*) is calculated by Pybel ^38^, a python wrapper of Openbabel ^39^. if the compound has ith fingerprint, Bit(i) = 1, else Bit(i) = 0. F_active_(*i*) and F_inactive_(*i*) are the frequency of ith fingerprint in the active and inactive compounds, respectively. α is the amplifying factor. In this study, α was optimized as 0.5 (**Figure S2**). Secondly, the Relative Frequency-Weighted Tanimoto Coefficient (RFW_TC) between two compounds was calculated as follows:

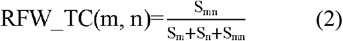

where RFW_TC(m,n) is RFW_TC between two compounds m and n. S_m_ and S_n_ are the sum of RFW_FPs in compound m and n, respectively. S_mn_ is the sum of the common RFW_FPs between two compounds. Finally, for each query chemical compounds, the maximum RFW_TC between the query and the active dataset (9584 compounds) was calculated. Then the P-value was calculated based on the maximum RFW_TC. As the maximum RFW_TC is less than 1.0 and the maximum RFW_TCs of the inactive compounds have a normal distribution (**Figure S3**), we can calculate the P-value as follows:

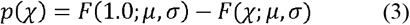

where p(χ) is the P-value at the maximum RFW_TC of x; F(χ;μ,σ) is the cumulative function of normal distribution. Using the maximum likelihood method ("fitdist" function in R "fitdistrplus" package ^40^), we estimated the location parameter μ of 0.461, the scale parameter σ of 0.121.

The SVM model was generated as follows. SVM is a powerful supervised learning algorithm suitable for non-linear classification problems ^41^. It is based on the idea of transforming data not linearly separable in feature space to a higher- or infinite-dimensional space where they can be separated linearly by a suitable soft-margin hyperplane ^42^. For our binary classification task, we firstly chosen kernel function and then performed a grid search of the penalty parameter C. The Scikit-learn Python wrappers for libsvm27 were used to choose kernel function and explore the hyper-parameter space ^43, 44^. The best-performing model was selected by plotting receiver operating characteristic (ROC) curve and calculating the area under the curve (AUC). The model with kernel function RBF and the penalty parameter C of 500 performed best (**Figure S4**). The detailed method for the selection of kernel function and the penalty parameter C can be found in **Part 5 of Supplemental material**.

The RF model was generated as follows. The algorithm of random forest is based on the ensemble of a large number of decision trees, where each tree gives a classification and the forest chooses the final classification having the most votes over all the trees in the forest ^45^. Random forest, implemented in Scikit-learn^43^, was used as a classifier with the following settings: (1) Number of trees was set to 900 (n_estimators =900). This parameter was selected by calculating AUC (**Figure S5**). (2) The minimum number of samples to split an internal node was set to 2 (min_samples_split = 2, default setting). (3) The minimum number of samples in newly created leaves was set to 1 (min_samples_leaf = 1, default setting). (4) The number of features to consider when looking for the best split was set to the square root of the number of descriptors (max_features = auto, default setting). (5) The maximum depth of the tree was expanded until all leaves are pure or until leaves contain less than min_samples_split samples (max_depth = none, default setting). (6) Bootstrap samples were used (bootstrap = true, default setting). For further documentation on the random forest implementation in Scikit-learn, the interested reader is referred to the website (http://scikit-learn.org).

### Performance evaluation of Anti-HIV-Predictor

To test the performance of Anti-HIV-Predictor, 10 runs of 5-fold cross-validation (CV) method (**Part 2 of Supplemental material**) were used to the three models (RFW_FP model, SVM model and

RF model). For each model, the ROC was plotted and the area under the curve (AUC) was calculated. The results of the CV tests were used to calculate the four quality indices: accuracy, precision, recall and F1 score. We used the default statistical definition for these quality indices:

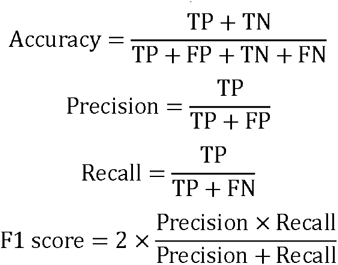

where true positive (TP) and true negative (TN) correspond to correctly predicted anti-HIV compound and non-anti-HIV, respectively, false positive (FP) denote non-anti-HIV compound predicted as anti-HIV compound, and false negative (FN) denote anti-HIV compound predicted as non-anti-HIV compound.

### Prediction of all compounds in ChEMBLdb using Anti-HIV-Predictor

All compounds with human target were downloaded from ChEMBLdb. A total of 251825 compounds and 1924 targets were obtained. They formed 384768 target-compound pairs. The anti-HIV activities of all compounds were predicted using **Anti-HIV-Predictor**. All compounds were screened rapidly by the three models. The cutoffs for RFW_TC model, SVM model and RF model are 0.05 (P-value), 0.5 (probability) and 0.5 (probability), respectively. The compound is predicted as anti-HIV compound if its anti-HIV activity supported by all three models (RFW_TC P-value < 0.05 and SVM probability > 0.5 and RF probability >0.5). Totally, 10488 compounds were predicted as anti-HIV compounds (**Supplemental Table S2**).

### Inferring the anti-HIV targets by enrichment analysis

After anti-HIV activity prediction of the 251825 compounds, we measured whether a target has the potential ability of anti-HIV using the APE method. The APE method is based on the results of anti-HIV activity prediction and uses a hypergeometric test to perform enrichment analysis. The P-value of each target can be calculated using the following equation:

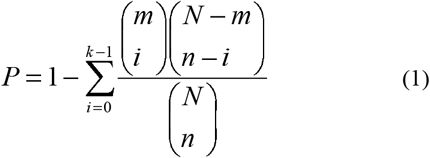

Here, N and n are the total number of compounds and the total number of anti-HIV compounds in ChEMBLdb, respectively; m and k represent the number of compounds and the number of anti-HIV compounds for a target, respectively. Both n and k are calculated above using Anti-HIV-Predictor. Because multiple tests (1924 targets) were performed, the Bonferroni correction method was used to adjust the P-value:

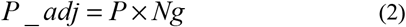

Here, P_adj is the adjusted P-value, P is the P-value of the hypergeometric test (without Bonferroni correction) and Ng is the number of human targets in the ChEMBLdb. Only targets with P_adj < 0.05 were considered as potential anti-HIV targets.

### Multiple features analysis of the potential anti-HIV targets

We studied these potential targets in details by analyzing multiple features including biology function, pathway, interaction, and druggability. The biology function and pathway of these targets were analyzed by GO and KEGG annotation in DAVID web server ^46^. The interactions of these targets were obtained by GeneMANIA analysis ^47,48^. The interactions of these targets with HIV proteins were obtained from NCBI HIV-1 human interaction database (**Supplemental Table S3**) ^23^. The network of interaction was visualized using Cytoscape v3.2 ^49^. The druggability of these targets was analyzed by comparing with the lists of approved targets and clinical targets (**Supplemental Table S4**) from DrugBank and therapeutic target database ^50-52^.

## Acknowledgements

This work was supported by the National Basic Research Program of China (Grant No. 2013CB835100), the National Natural Science Foundation of China (Grant Nos. 31401142 and 31401137) and the Instruments Function Deployment Foundation of CAS (2014gk01).

## Author contribution

S-X D and J-F H conceived and designed the research. S-X D, W-X L, H-J L, J-Q L, J-J Z, Q W, B-WC, Y-D G and G-H L performed data analysis. S-X D, W-X L, H-J L, G-H L and J-F H wrote or contributed to the writing of the manuscript. All authors reviewed the manuscript.

## References

1 O'Brien, S. J. & Hendrickson, S. L. Host genomic influences on HIV/AIDS. Genome biology 14, 201, doi:10.1186/gb-2013-14-1-201 (2013).

2 Menendez-Arias, L. Molecular basis of human immunodeficiency virus type 1 drug resistance: overview and recent developments.Antiviral research 98, 93–120, doi:10.1016/j.antiviral.2013.01.007 (2013).

3 Sliva, K. Latest animal models for anti-HIV drug discovery. Expert opinion on drug discovery 10, 111–123, doi:10.1517/17460441.2015.975201 (2015).

4 De Clercq, E. Anti-HIV drugs: 25 compounds approved within 25 years after the discovery of HIV. International journal of antimicrobial agents 33, 307–320, doi:10.1016/j.ijantimicag.2008.10.010 (2009).

5 Swaminathan, G., Navas-Martin, S. & Martin-Garcia, J. MicroRNAs and HIV-1 infection: antiviral activities and beyond. Journal of molecular biology426, 1178–1197, doi:10.1016/j.jmb.2013.12.017 (2014).

6 Zhan, P., Pannecouque, C., De Clercq, E. & Liu, X. Anti-HIV Drug Discovery and Development: Current Innovations and Future Trends. Journal of medicinal chemistry 59, 2849–2878, doi:10.1021/acs.jmedchem.5b00497 (2016).

7 Chabria, S. B., Gupta, S. & Kozal, M. J. Deep sequencing of HIV: clinical and research applications. Annual review of genomics and human genetics 15, 295–325, doi:10.1146/annurev-genom-091212-153406 (2014).

8 Law, G. L., Korth, M. J., Benecke, A. G. & Katze, M. G. Systems virology: host-directed approaches to viral pathogenesis and drug targeting. Nature reviews. Microbiology 11, 455–466, doi:10.1038/nrmicro3036 (2013).

9 Shea, P. R., Shianna, K. V., Carrington, M. & Goldstein, D. B. Host genetics of HIV acquisition and viral control. Annual review of medicine 64, 203–217, doi:10.1146/annurev-med-052511-135400 (2013).

10 Ballana, E. & Esté, J. A. Insights from host genomics into HIV infection and disease: identification of host targets for drug development. Antiviral research 100, 473–486 (2013).

11 Law, G. L., Tisoncik-Go, J., Korth, M. J. & Katze, M. G. Drug repurposing: a better approach for infectious disease drug discovery? Current opinion in immunology 25, 588–592, doi:10.1016/j.coi.2013.08.004 (2013).

12 Li, B. Q. et al. Identifying chemicals with potential therapy of HIV based on protein-protein and protein-chemical interaction network. PloS one 8, e65207, doi:10.1371/journal.pone.0065207 (2013).

13 Ma-Lauer, Y., Lei, J., Hilgenfeld, R. & von Brunn, A. Virus-host interactomes--antiviral drug discovery. Current opinion in virology 2, 614–621, doi:10.1016/j.coviro.2012.09.003 (2012).

14 Coley, W., Kehn-Hall, K., Duyne, R. V. & Kashanchi, F. Novel HIV-1 therapeutics through targeting altered host cell pathways. Expert opinion on biological therapy 9, 1369–1382 (2009).

15 Amaya, M. et al. Proteomic strategies for the discovery of novel diagnostic and therapeutic targets for infectious diseases. Pathogens and disease 71, 177–189, doi:10.1111/2049-632X.12150 (2014).

16 Zhou, H. et al. Genome-scale RNAi screen for host factors required for HIV replication. Cell host & microbe 4, 495–504, doi:10.1016/j.chom.2008.10.004 (2008).

17 Jaeger, S. et al. Global landscape of HIV-human protein complexes. Nature 481, 365–370 (2012).

18 König, R. et al. Global analysis of host-pathogen interactions that regulate early-stage HIV-1 replication. Cell 135, 49–60 (2008).

19 Park, R. J. et al. A genome-wide CRISPR screen identifies a restricted set of HIV host dependency factors. Nature genetics 49, 193–203 (2017).

20 Griffith, M. et al. DGIdb: mining the druggable genome. Nature methods 10, 1209–1210, doi:10.1038/nmeth.2689 (2013).

21 Russ, A. P. & Lampel, S. The druggable genome: an update. Drug discovery today 10, 1607–1610, doi:10.1016/S1359-6446(05)03666-4 (2005).

22 Hopkins, A. L. & Groom, C. R. The druggable genome. Nature reviews. Drug discovery 1, 727–730, doi:10.1038/nrd892 (2002).

23 Ako-Adjei, D. et al. HIV-1, human interaction database: current status and new features. Nucleic acids research 43, D566–570, doi:10.1093/nar/gku1126 (2015).

24 Li, G.-H. & Huang, J.-F. Inferring therapeutic targets from heterogeneous data: HKDC1 is a novel potential therapeutic target for cancer. Bioinformatics 30, 748–752 (2014).

25 Kim, J. H., Excler, J. L. & Michael, N. L. Lessons from the RV144 Thai phase III HIV-1 vaccine trial and the search for correlates of protection. Annual review of medicine 66, 423–437, doi:10.1146/annurev-med-052912-123749 (2015).

26 Sadanand, S., Suscovich, T. J. & Alter, G. Broadly Neutralizing Antibodies Against HIV: New Insights to Inform Vaccine Design. Annual review of medicine 67, 185–200, doi:10.1146/annurev-med-091014-090749 (2016).

27 A Waheed, A. & Tachedjian, G. Why Do We Need New Drug Classes for HIV Treatment and Prevention? Current topics in medicinal chemistry 16, 1343–1349 (2016).

28 Bui, Q. C., Nuallain, B. O., Boucher, C. A. & Sloot, P. M. Extracting causal relations on HIV drug resistance from literature. BMC bioinformatics 11, 101, doi:10.1186/1471-2105-11-101 (2010).

29 Beerenwinkel, N. et al. Computational methods for the design of effective therapies against drug resistant HIV strains. Bioinformatics 21, 3943–3950, doi:10.1093/bioinformatics/bti654 (2005).

30 Alkhatib, G. The biology of CCR5 and CXCR4. Current opinion in HIV and AIDS 4, 96–103, doi:10.1097/COH.0b013e328324bbec (2009).

31 Murphy, P. M. Viral exploitation and subversion of the immune system through chemokine mimicry. Nature immunology 2, 116–122 (2001).

32 Sodhi, A., Montaner, S. & Gutkind, J. S. Viral hijacking of G-protein-coupled-receptor signalling networks. Nature Reviews Molecular Cell Biology 5, 998–1012 (2004).

33 Gaulton, A. et al. ChEMBL: a large-scale bioactivity database for drug discovery. Nucleic acids research 40, D1100–1107, doi:10.1093/nar/gkr777 (2012).

34 Gaulton, A. et al. The ChEMBL database in 2017. Nucleic acids research 45, D945–D954, doi:10.1093/nar/gkw1074 (2017).

35 Sausville, E. A. & Shoemaker, R. H. Role of the National Cancer Institute in acquired immunodeficiency syndrome-related drug discovery. Journal of the National Cancer Institute. Monographs, 55–57 (2001).

36 Li, G. H. & Huang, J. F. CDRUG: a web server for predicting anticancer activity of chemical compounds. Bioinformatics 28, 3334–3335, doi:10.1093/bioinformatics/bts625 (2012).

37 Dai, S. X. et al. In silico identification of anti-cancer compounds and plants from traditional Chinese medicine database. Scientific reports 6, 25462, doi:10.1038/srep25462 (2016).

38 O'Boyle, N. M., Morley, C. & Hutchison, G. R. Pybel: a Python wrapper for the OpenBabel cheminformatics toolkit. Chemistry Central journal 2, 5, doi:10.1186/1752-153X-2-5 (2008).

39 O'Boyle, N. M. et al. Open Babel: An open chemical toolbox. Journal of cheminformatics 3, 33, doi:10.1186/1758-2946-3-33 (2011).

40 Delignette-Muller, M. L. & Dutang, C. fitdistrplus: An R Package for Fitting Distributions. J. Stat. Softw 64, 1–34 (2014).

41 Orru, G., Pettersson-Yeo, W., Marquand, A. F., Sartori, G. & Mechelli, A. Using Support Vector Machine to identify imaging biomarkers of neurological and psychiatric disease: a critical review. Neuroscience and biobehavioral reviews 36, 1140–1152, doi:10.1016/j.neubiorev.2012.01.004 (2012).

42 Burges, C. J. C. A tutorial on Support Vector Machines for pattern recognition. Data Min Knowl Disc 2, 121–167, doi:Doi 10.1023/A:1009715923555 (1998).

43 Pedregosa, F. et al. Scikit-learn: Machine Learning in Python. J Mach Learn Res 12, 2825–2830 (2011).

44 Chang, C. C. & Lin, C. J. LIBSVM: A Library for Support Vector Machines. Acm T Intel Syst Tec 2, doi:Artn 2710.1145/1961189.1961199 (2011).

45 Breiman, L. Random forests. Mach Learn 45, 5–32, doi:Doi 10.1023/A:1010933404324 (2001).

46 Huang, D. W., Sherman, B. T. & Lempicki, R. A. Systematic and integrative analysis of large gene lists using DAVID bioinformatics resources. Nature protocols 4, 44–57 (2009).

47 Montojo, J., Zuberi, K., Rodriguez, H., Bader, G. D. & Morris, Q. GeneMANIA: Fast gene network construction and function prediction for Cytoscape. F1000Research 3, 153, doi:10.12688/f1000research.4572.1 (2014).

48 Warde-Farley, D. et al. The GeneMANIA prediction server: biological network integration for gene prioritization and predicting gene function. Nucleic acids research 38, W214–220, doi:10.1093/nar/gkq537 (2010).

49 Shannon, P. et al. Cytoscape: a software environment for integrated models of biomolecular interaction networks. Genome research 13, 2498–2504, doi:10.1101/gr.1239303 (2003).

50 Yang, H. et al. Therapeutic target database update 2016: enriched resource for bench to clinical drug target and targeted pathway information. Nucleic acids research 44, D1069–1074, doi:10.1093/nar/gkv1230 (2016).

51 Chen, X., Ji, Z. L. & Chen, Y. Z. TTD: Therapeutic Target Database. Nucleic acids research 30, 412–415 (2002).

52 Wishart, D. S. et al. DrugBank: a knowledgebase for drugs, drug actions and drug targets. Nucleic acids research 36, D901–906, doi:10.1093/nar/gkm958 (2008).

